# Hybridization Capture Using RAD Probes (hyRAD), a New Tool for Performing Genomic Analyses on Collection Specimens

**DOI:** 10.1101/025551

**Authors:** Tomasz Suchan, Camille Pitteloud, Nadezhda S. Gerasimova, Anna Kostikova, Sarah Schmid, Nils Arrigo, Mila Pajkovic, Michał Ronikier, Nadir Alvarez

## Abstract

In the recent years, many protocols aimed at reproducibly sequencing reduced-genome subsets in non-model organisms have been published. Among them, RAD-sequencing is one of the most widely used. It relies on digesting DNA with specific restriction enzymes and performing size selection on the resulting fragments. Despite its acknowledged utility, this method is of limited use with degraded DNA samples, such as those isolated from museum specimens, as these samples are less likely to harbor fragments long enough to comprise two restriction sites making possible ligation of the adapter sequences (in the case of double-digest RAD) or performing size selection of the resulting fragments (in the case of single-digest RAD). Here, we address these limitations by presenting a novel method called hybridization RAD (hyRAD). In this approach, biotinylated RAD fragments, covering a random fraction of the genome, are used as baits for capturing homologous fragments from genomic shotgun sequencing libraries. This simple and cost-effective approach allows sequencing of orthologous loci even from highly degraded DNA samples, opening new avenues of research in the field of museum genomics. Not relying on the restriction site presence, it improves among-sample loci coverage. In a trial study, hyRAD allowed us to obtain a large set of orthologous loci from fresh and museum samples from a non-model butterfly species, with a high proportion of single nucleotide polymorphisms present in all eight analyzed specimens, including 58-year-old museum samples. The utility of the method was further validated using 49 museum and fresh samples of a Palearctic grasshopper species for which the spatial genetic structure was previously assessed using mtDNA amplicons. The application of the method is eventually discussed in a wider context. As it does not rely on the restriction site presence, it is therefore not sensitive to among-sample loci polymorphisms in the restriction sites that usually causes loci dropout. This should enable the application of hyRAD to analyses at broader evolutionary scales.

## Introduction

With the advent of next-generation sequencing, conducting genomic-scale studies on non-model species has become a reality [1]. The cost of genome sequencing has substantially dropped over the last decade and repositories now encompass an incredible amount of genomic data, which has opened avenues for the emerging field of ecological genomics. However, when working at the population level—at least in eukaryotes—sequencing whole genomes still lies beyond the capacities of most laboratories, and a number of techniques targeting a subset of the genome have been developed [2, 3]. Among the most popular are approaches relying on hybridization capture of exome [4] or conserved fragments of the genome [5], RNA sequencing (RNAseq [6]), and Restriction-Associated-DNA sequencing (RADseq [7, 8]). The latter has been developed in many different versions, but generally relies on specific enzymatic digestion and further selection of a range of DNA fragment sizes. RAD-sequencing has proved to be a cost-and time-effective method of SNP (single nucleotide polymorphisms) discovery, and currently represents the best tool available to tackle questions in the field of molecular ecology. The wide utility of RAD-sequencing in ecological, phylogenetic and phylogeographic studies is however limited by two main factors: i) the quality of the starting genomic DNA; ii) the degree of divergence among the studied specimens, that translates into DNA sequence polymorphism at the restriction sites targeted by the RAD protocols.

Sequence polymorphism at the DNA restriction site causes a progressive loss of shared restriction sites among diverging clades and results in null alleles for which sequence data cannot be obtained. This limitation critically reduces the number of orthologous loci that can be surveyed across the complete set of analyzed specimens and leads to biased genetic diversity estimates [9-11]. This phenomenon, combined with other technical issues – such as polymerase chain reaction (PCR) competition effects – is a serious limitation of most classic RAD-sequencing protocols that needs to be addressed.

In addition, RAD-sequencing protocols rely on relatively high molecular weight DNA (especially for the ddRAD protocol [12]), notably because enzyme digestion and size selection of resulting fragments are the key steps for retrieving sequence data across orthologous loci. Therefore, RAD-sequencing cannot be applied to degraded DNA samples, a limitation also shared by classical genotyping methods and amplicon sequencing. Museum collections, although encompassing samples covering large spatial areas and broad temporal scales, have not necessarily ensured optimal conditions for DNA preservation. As a result, many museum specimens yield highly fragmented DNA – even for relatively recently collected samples [13-15], limiting their use for molecular ecology, conservation genetics, phylogeographic and phylogenetic studies [16, 17]. A cost-effective and widely applied approach for genomic analyses on museum specimens would allow exploring often unique biological collections, e.g., encompassing rare or now extinct taxa/lineages or organisms occurring ephemerally in natural habitats and posing problems for sampling. It would also allow studying temporal shifts in genetic diversity using historical collections, now applied only in a handful of cases at a genomic scale [18].

Hybridization-capture methods have been acknowledged as a promising way to address both the allele representation and DNA quality limitations [19, 20]. Such approaches however usually rely on prior genome/transcriptome knowledge and until recently have been largely confined to model organisms. Addressing this limitation, the recent development of UltraConserved Elements (UCE [3, 5]) or anchored hybrid enrichment [21] capture-based methods allowed targeting homologous loci at broad phylogenetic scales using one set of probes. It however requires a time-consuming design and costly synthesis of the probes for capturing the DNA sequences of interest. Similarly, exon capture techniques, recently applied in the field of museum genomics [18], require fresh specimens for RNA extraction or synthesizing the probes based on the known transcriptome.

Here, we present an approach we called ‘hybridization RAD’ (hyRAD), in which DNA fragments, generated using double digestion RAD protocol (ddRAD [8]) applied to fresh samples, are used as hybridization-capture probes to enrich shotgun libraries in the fragments of interest. Our method thus combines the simplicity and relatively low cost of developing RAD-sequencing libraries with the power and accuracy of hybridization-capture methods. This enables the effective use of low quality DNA and limits the problems caused by sequence polymorphisms at the restriction site. Moreover, utilizing standard ddRAD and shotgun sequencing protocols allows application of the hyRAD protocol in laboratories already utilizing the abovementioned methods, for little cost.

In short, the hyRAD approach consists of the following steps (Fig 1):

1. generation of a ddRAD library based on high-quality DNA samples, narrow size selection of the resulting fragments and removing adapter sequences;
2. biotinylation of the resulting fragments, hereafter called the probes;
3. construction of a shotgun sequencing library from DNA samples (either fresh or degraded as in museum specimens);
4. hybridization capture of the resulting shotgun libraries on the probes;
5. sequencing of enriched shotgun libraries and optionally of the ddRAD library (probes precursor) for further use as a reference;
6. bioinformatic treatment (Fig 2): assembly of the reads into contigs, alignment to the sequenced ddRAD library or *de novo* assembly, SNP calling.

**Fig 1.**
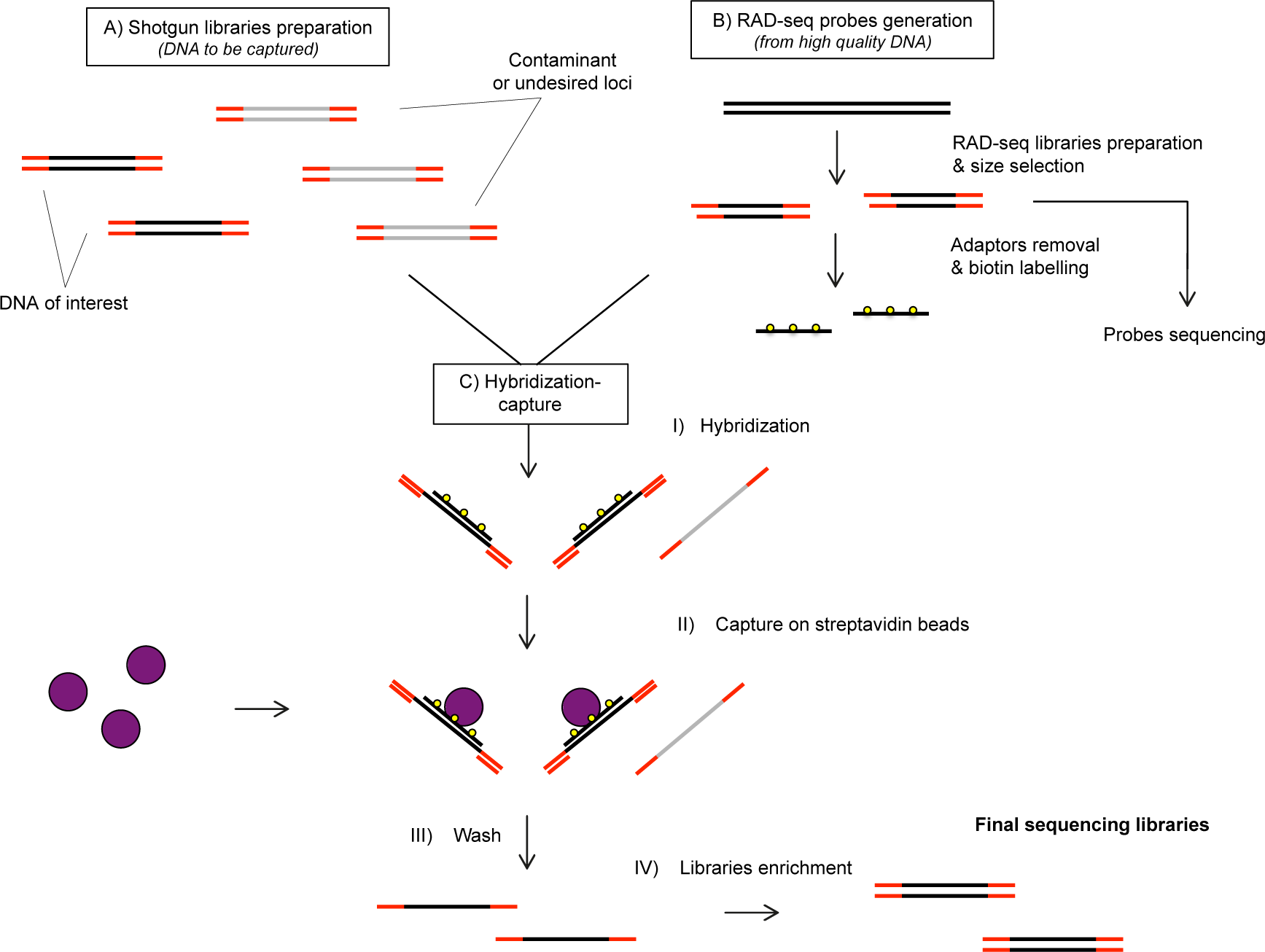
Lab-work procedure used for hyRAD development. Homologous reads from shotgun genomic libraries are captured through hybridization on random RAD-based probes. These fragments are then separated using streptavidin-coated beads and sequenced.

In this paper, we describe the laboratory and bioinformatic pipelines for obtaining hyRAD data, as well as validate the usefulness of the method on two empirical datasets. We first test the method on the DNA obtained from museum and fresh specimens of *Lycaena helle* butterflies. We explore different bioinformatic approaches for assembling loci out of the hybridization-capture libraries, namely: i) mapping captured libraries on previously sequenced RAD loci from fresh samples; ii) using RAD loci as seeds and the captured libraries’ reads to extend the RAD loci in order to obtain longer loci for mapping; iii) *de novo* assembly of the captured reads from a single, well-preserved butterfly specimen for the reference. Secondly, as a proof of concept, we apply the protocol to museum and fresh samples of *Oedaleus decorus*, a Palearctic grasshopper species for which a marked east-west spatial genetic structure has been identified in a previous study [22].

## Materials and Methods

### Study species and study design

For the first step of the method development, we used samples of the butterfly *Lycaena helle* (Lepidoptera, Lycaenidae) (Table 1). Three recently collected, ethanol-preserved samples from Romania, France and Kazakhstan were used for generating the RAD probes, in order to cover variation within the full species range. Genomic libraries to be enriched by sequence capture were built using eight samples which included seven museum dry-pinned specimens from Finland (4 collected in 1985 and 3 collected in 1957) and one recently collected and ethanol-preserved specimen from Romania. Using these eight samples we compared the outputs between fresh and historical DNA of different age, and tested the importance of DNA sonication in each case. The museum samples were loaned from the Finnish Museum of Natural History in Helsinki, and the ethanol-preserved samples were obtained from Roger Vila’s Butterfly Diversity and Evolution lab (Institute of Evolutionary Biology, CSIC, Barcelona, Spain).

**Table 1.** *Lycaena helle* samples used in the study.

| Type of preservation | Year of collection | DNA concentration [ng/ul] | Locality |
| --- | --- | --- | --- |
| dry (pin-mounted) | 1957 | 2.12 | Kuusamo, Finland |
| dry (pin-mounted) | 1957 | 3.02 | Kuusamo, Finland |
| dry (pin-mounted) | 1957 | 1.48 | Kuusamo, Finland |
| dry (pin-mounted) | 1985 | 29.2 | Kuusamo, Finland |
| dry (pin-mounted) | 1985 | 19.7 | Kuusamo, Finland |
| dry (pin-mounted) | 1985 | 17.5 | Kuusamo, Finland |
| dry (pin-mounted) | 1985 | 8.84 | Kuusamo, Finland |
| ethanol | 2007 | 2.12 | Dumbrava Vadului, Romania |

For the method validation, we used 53 samples of the grasshopper *Oedaleus decorus*, including 49 samples of both fresh and museum collection specimens for constructing genomic libraries for the capture (see Table 2), and four fresh specimens spanning the species’ distribution (Switzerland, Spain, Hungary, Russia) for generating the RAD probes. The museum samples were on average 64-years-old, with the oldest sample dating back to 1908 and were provided by the National History Museum of London (UK), the Natural History Museum of Bern (Switzerland), the Zoological Museum of Lausanne (Switzerland), the ETH Entomological Collection (Switzerland), the Natural History Museum of Basel (Switzerland), the Natural History Museum of Geneva (Switzerland), and the Natural History Museum of Zurich (Switzerland). The four fresh grasshopper samples were provided by G. Heckel (University of Bern).

**Table 2.** Summary of *Oedaleus decorus* samples used in the study.

| Type of preservation | Mean year of collection (range) | Mean DNA concentration [ng/ul] ± SD (range) | Localities |
| --- | --- | --- | --- |
| ethanol | 2006 (2005-2009) | 28.34 ± 28.04 (3.2-105.7) | Croatia, France, Italy, Russia, Spain, Switzerland |
| dry (pin-mounted) | 1952 (1908-1997) | 18.31 ± 21.73 (0.3-121.4) | Algeria, France, Greece, Italy, Madeira,, Portugal (mainland and Madeira), Spain (mainland and Canary islands), Switzerland, Turkey |

### DNA extraction

DNA was extracted from insect legs for all the samples. As museum specimens are usually characterized by low-content of degraded DNA, the isolation protocol was optimized accordingly. The samples were extracted using QIAamp DNA Micro kit (Qiagen, Hombrechtikon, Switzerland) in a laboratory dedicated to low-DNA content samples at the University of Lausanne, Switzerland. For these samples, DNA recovery was improved by prolonged sample grinding, overnight incubation in the lysis buffer for 14h and final DNA elution in 20 μl of the buffer with gradual column centrifugation. Extraction of fresh samples was performed using DNeasy Blood & Tissue Kit (Qiagen). DNA extraction and library preparation using museum specimens was performed using consumables dedicated to the museum specimens only. Benches were thoroughly cleaned with bleach and filter tips were used at all stages of lab work.

### RAD probes preparation

The probe precursors were prepared using a double-digestion RAD protocol [8, 23], with further modifications.

Total genomic DNA was digested at 37°C for 3 hours in a 9 μl reaction, containing 6 μl of DNA, 1x CutSmart buffer (New England Biololabs – NEB, Ipswich, MA, USA), 1 U MseI (NEB) and 2 U of SbfI-HF (NEB). The reaction products were purified using AMPure XP (Beckman Coulter, Brea, USA), with a ratio of 2:1 with the sample, according to the manufacturer’s instructions, and resuspended in 10 μl of 10 mM Tris buffer. Subsequently, adapters were ligated to the purified restriction-digested DNA in a 20 μl reaction containing 10 μl of the insert, 0.5 μM of RAD-P1 adapter, 0.5 μM of universal RAD-P2 adapter, 1x T4 ligase buffer, and 400 U of T4 DNA ligase (NEB). Adapter sequences are shown in Table 3; single strand adapter oligonucleotides are annealed before use by heating to 95°C and gradual cooling. Ligation was performed at 16°C for 3 hours. The reaction products were purified using an AMPure XP ratio 1:1 with the sample, and resuspended in 10 mM Tris buffer. The ligation product was size-selected using the Pippin Prep electrophoresis platform (Sage Science, Beverly, USA) with a peak at 270 bp and ‘tight’ size selection range.

**Table 3.**
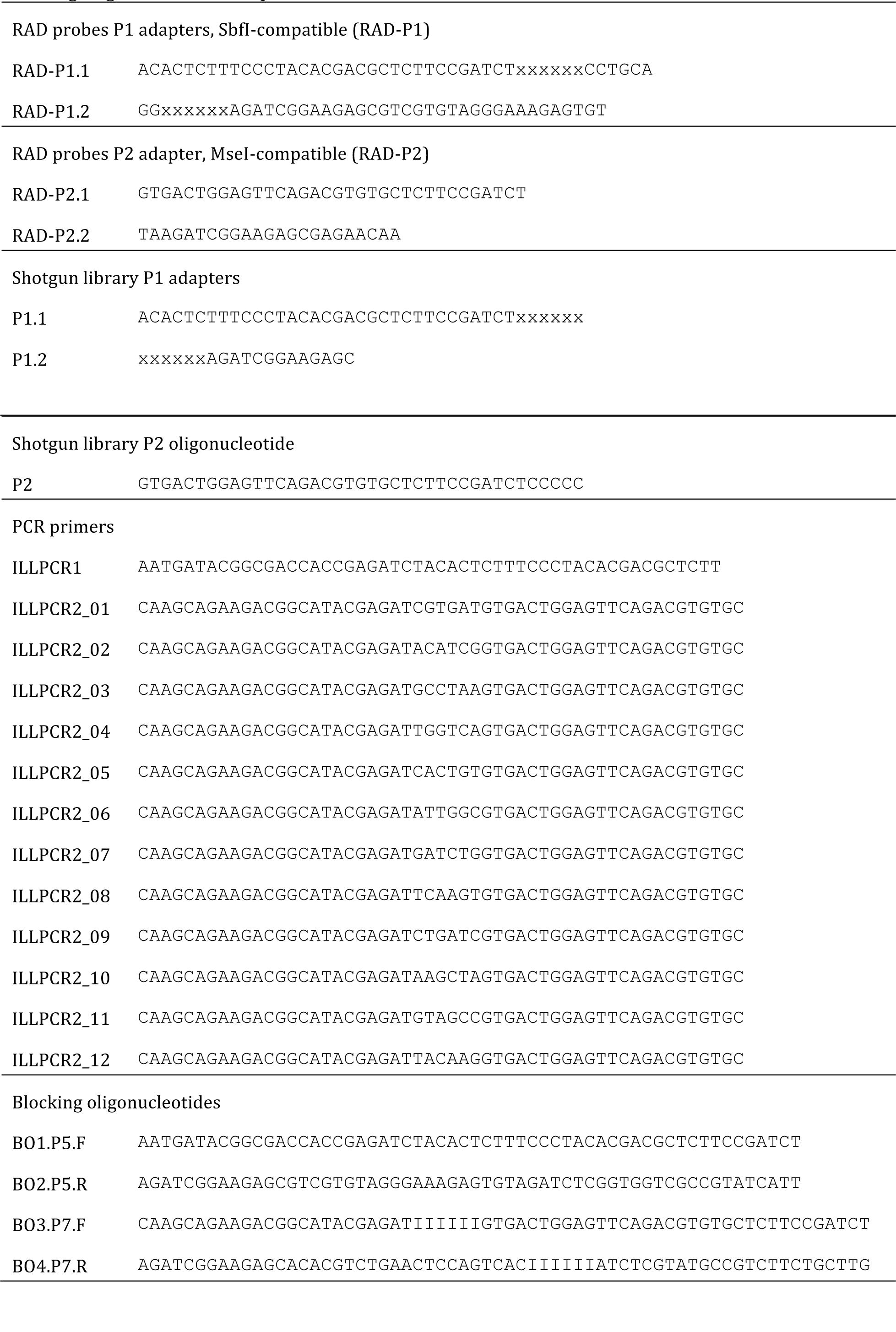
Oligonucleotides used in the protocol. x = barcode sequence in the adapters; barcode sequences can be designed using published scripts [24], available at: https://bioinf.eva.mpg.de/multiplex/; I = inosine in the region complementary to the barcode in blocking oligonucleotides sequences.

The resulting template was amplified by PCR in a 10 μl mix consisting of 1x Q5 buffer, 0.2 mM of each dNTP, 0.6 μM of each primer (Table 3), and 0.2 U Q5 hot-start polymerase (NEB). The thermocycler program included initial denaturation for 30 sec at 98°C; 30 PCR cycles of 20 sec at 98°C, 30 sec at 60°C, and 40 sec at 72°C; followed by a final extension for 10 min at 72°C. In order to obtain sufficient amount of material for the probes, the reaction had to be run in replicates. The necessary number of replicates has to be determined empirically to reach in total 500-1000 ng of the amplified product required for each capture. DNA amounts were assessed using a Qubit fluorometer (Waltham, MA, USA). Success of size selection and of PCR reactions was confirmed by running them on a Fragment Analyzer (see S1 Fig; Advanced Analytical, Ankeny, IA, USA; see S1 Fig). Afterwards, the PCR products were pooled and purified using AMPure XP, with a ratio of 1:1 with the amplified DNA volume.

An aliquot of the resulting library was sequenced and the rest of the library was converted into probes by removing adapter sequences by enzymatic restriction followed by biotinylation. The probe precursors were incubated at 37°C for 3 hours in a 50 μl reaction containing 30 ¼l of DNA, 1x CutSmart buffer (NEB), 5 U of MseI (NEB) and 10 U of SbfI-HF (NEB), replicated as required by the amount of the amplified product. The reaction was ended with 20 min enzyme inactivation at 65°C for 20 min and the resulting fragments were purified using AMPure XP:reaction volume ratio of 1.5:1. Purified fragments were biotin nick-labelled using BioNick DNA Labeling System (Thermo Fisher, Waltham, MA, USA) according to the supplier’s instructions and purified using AMPure XP:reaction volume ratio of 1.5:1. The resulting fragments will thereafter be referred to as probes.

### Shotgun library preparation

Shotgun libraries were prepared from the fresh and museum specimens based on a published protocol for degraded DNA samples [15], modified in order to incorporate adapter design of Meyer & Kircher [24]. The approach used for library preparation, utilizing barcoded P1 adapter and 12 indexed P2 PCR primers, allows a high sample multiplexing on a single sequencing lane (see Table 3 [24]).

For *L. helle*, DNA from each individual was divided in two aliquots. One aliquot was kept intact (i.e. high molecular weight DNA in the fresh sample and naturally degraded DNA in museum specimens) and the second was sonicated using Covaris focused ultrasonicator (Woburn, MA, USA) with a peak at 300 bp. Both aliquots were processed in parallel during subsequent steps of libraries preparation. All the libraries from *O. decorus* were prepared without sonication, based on the test results obtained from the *L. helle* libraries.

DNA samples were first 5’-phosphorylated in order to allow adapter ligation in the next steps of the protocol. 8 μl of DNA was denatured at 95°C for 10 minutes and quickly chilled on ice. The 10 μl reaction consisting of denatured DNA, 1x PNK buffer and 10U of T4 polynucleotide kinase (NEB) was incubated at 37°C for 30min and heat-inactivated at 65°C for 20 min. The DNA was then purified using an AMPure:reaction volume ratio of 2:1 and resuspended in 10 μl of 10 mM Tris buffer.

A guanidine tailing reaction of the 3’-terminus was performed after heat denaturation ofDNA at 95°C for 10 minutes and quickly chilling on ice. The reaction composed of 1x buffer 4 (NEB), 0.25 mM cobalt chloride (NEB), 4 mM GTP (Life Technologies), 10 U TdT (NEB) and 10 μl of denatured DNA in 20 μl reaction volume was incubated at 37°C for 30 min and heat-inactivated at 70°C for 10 min.

The second DNA strand was synthesized using Klenow Fragment (3’ → 5’ exo-), with a primer consisting of the Illumina P2 sequence and a poly-C sequence homologous to the poly-G tail (see Table 3) added to the DNA strand in the previous reaction. A 10 μl reaction mix consisting of 1 μl of NEBuffer 4 (10x), 0.6 μl of dNTP mix (25 mM each), 1 μlof the P2 oligonucleotide (15 mM), 5.4 μl of water, and 2 μl of Klenow Fragment (’ — 5’ exo-; NEB, 5 U/μl) was added to the 20 μl of the TdT reaction mix, incubated at 23°C for 3 h, and heat-inactivated at 75°C for 20 min. The double stranded product was then blunt-ended by adding a mix consisting of 0.5 μl of NEBuffer 4 (10x), 0.35 μl of BSA (10 mg/ml), 0.2 μ of T4 DNA polymerase (NEB, 3 U/μl) and 3.95 μl of water, and incubated at 12°C for 15 min. The resulting product was purified using AMPure XP:reaction ratio of 2:1 and resuspended in 10 μl of 10 mM Tris buffer.

Barcoded P1 adapters (see Table 3) were ligated to the 5’-phosphorylated end of the double-stranded product in a 20 μl reaction consisting of 10 μl of the double-stranded DNA, 1 μl of the 25 uM adapters, 1x T4 DNA ligase buffer, and 400 U of T4 DNA ligase (NEB). Adapters have to be annealed before use in the RAD probes protocol. The reaction was incubated at 16°C overnight. The resulting product was purified using an AMPure:reaction ratio of 1:1 and resuspended in 20 μl of 10 mM Tris buffer. Ligated P1 adapters were filled-in in a 40 μl reaction consisting of 20 μl of purified ligation product, 1x ThermoPol reaction buffer (NEB), 12 U of Bst polymerase (NEB), and dNTPs (0.25 mM each), and incubated at 37°C for 20 min.

The resulting template was amplified by PCR adding 15 μl of a mix consisting of 5 μl of Q5 reaction buffer (5x), 0.2 μl of dNTPs (25 mM each), 2.5 μl of the PCR primer mix (5 μM each), and 0.5 U of Q5 Hot Start High-Fidelity DNA polymerase (NEB) to the 10 μl of the template. The program started with 20 sec at 98°C, followed by 25 cycles of 10 sec at 98°C, 20 sec at 60°C, and 25 sec at 72°C, followed by a final extension for 2 min at 72°C. Success of each PCR reaction was checked using gel electrophoresis, and the resulting products were purified using AMPure XP:reaction ratio of 0.7:1. Samples were then pooled in equimolar ratios.

### In solution hybridization capture, library reamplification and sequencing

The hybridization capture and library enrichment steps described below are based on previously published protocols [13, 25] with some modifications. The hybridization mix consisted of 6x SSC, 50 mM EDTA, 1% SDS, 2x Denhardt’s solution, 2 μM of each blocking oligonucleotide (to prevent hybridization of adapter sequences; see (Table 3), 500 to 1000 ng of the probes and 500 to 1000 ng of the shotgun libraries, in a total volume of 40 μΙ On account of grasshopper larger genome size and preliminary results indicating low signal-noise ratio in the butterfly libraries, 500 ng of human Cot-1 DNA (Thermo Fisher Scientific, Switzerland) was added to the *O. decorus* hybridization mix in order to prevent non-specific hybridizations caused by repetitives equences. The mix was denatured at 95°C for 10 min and subsequently incubated at 65°C for 48 hours. The probes, hybridized with targeted fragments of the library, were then separated on streptavidin beads (Dynabeads M-280, Life Technologies). 10 μl of the beads solution was washed three times on the magnet with 200 μl of TEN buffer (10 mM Tris-HCl 7.5, 1 mM EDTA, 1 M NaCl) and resuspended in 200 μl of TEN. 40 μl of the hybridization mix was added to the 200 μl of the beads solution and incubated for 30 min at room temperature. After separating the beads with the magnet, the supernatant was removed and the beads were washed four times under different stringency conditions as follows. The beads were resuspended in 200 μl of 65°C 1x SSC/0.1% SDS wash buffer, incubated for 15 min at 65°C, separated on the magnet and the supernatant was removed. The above step was performed again with 1x SSC/0.1% SDS, followed by 0.5x SSC/0.1% SDS and 0.1x SSC/0.1% SDS. Finally, the hybridization-enriched product was washed-off from the probes by adding 30 μl of 80°C water and incubating at 80°C for 10 min.

Enrichment of the captured libraries was performed in a 50 μl PCR reaction containing 1xQ5 reaction buffer (NEB), 0.2 mM dNTPs, 0.5 μM of each PCR primer (the P1 universal primer and one of the 12 P2 indexed primers, see (Table 3), 1U of Q5 Hot Start High-Fidelity DNA Polymerase (NEB), and 15 μl of the template. The program started with 20 sec initial denaturation at98°C; followed by 25 PCR cycles of 10 sec at 98°C, 20 sec at 60°C, and 25 sec at 72°C; and a final extension for 2 min at 72°C. The enriched-captured libraries were purified using an AMPure XP:reaction ratio 1:1 and pooled in equimolar ratios for sequencing (see S2 Fig for a profile example of the re-amplified capture library after AMPure purification).

The probes precursors (RAD library) for the butterfly libraries were sequenced on one lane of Illumina MiSeq 300 bp single-end. Butterfly capture-enriched libraries were sequenced on one lane of MiSeq 150 bp paired-end, and grasshopper capture-enriched libraries were sequenced on one lane of Illumina HiSeq 100 bp paired-end.

Updated versions of the lab protocol can be found at https://github.com/chiasto/hyRAD.

### Data analysis

The hyRAD datasets correspond to target-enriched libraries and cannot be analysed with the usual RAD pipelines [26, 27] Indeed, although they were generated using RAD loci, the obtained sequences are not flanked by the restriction sites and instead may not overlap completely and/or extend before and after the RAD locus. As a result, the analysis pipeline must include the following steps:

1. demultiplexing and cleaning of raw reads;
2. building of reference sequences for each RAD locus;
3. alignment of reads against the obtained references;
4. SNP calling.

All bioinformatic steps of the hyRAD pipeline can be run at https://insidedna.me.

### Demultiplexing and data preparation

The obtained reads were demultiplexed using the <u>fastx barcode splitter tool</u> from the <u>FASTX-Toolkit</u> package [28]. RAD-seq sequences (probe precursors) were processed by Trim Galore! [29] and cleaned with the <u>fastq-mcf tool</u> from ea-utils package [30] to remove low quality nucleotides and adapter sequences. The PCR duplicates were removed from RAD-seq probes precursors and hyRAD datasets using the MarkDuplicates tool of Picard toolkit [31]. Reads from hyRAD libraries were tested for exogeneous DNA contamination using BLAST against NCBI nucleotide database (50,000 reads for sonicated or non-sonicated fresh or museum DNA samples).

### Exploring the methods of reference creation on *Lycaena helle* libraries

Paired-end reads obtained from the hybridization-capture library for each sample were mapped onto three references: (1) consensus sequences for the clustered RAD-seq reads (RAD-ref), (2) RAD-seq reads extended using hybridization-captured reads (RAD-ref-ext), and (3) contigs assembled from the reads of hybridization-captured samples (assembly-ref). As we had no reference genome available, we focused on checking the numbers of loci/SNPs obtained by mapping the reads on each reference and the overlap of the loci obtained using different methods. Theoretically, we should be able to retrieve all the loci from the sequenced probe precursors (RAD-seq library) in the captured libraries. We thus evaluate the number of captured loci homologous to those retrieved by sequencing the RAD-seq libraries.However, different factors affect signal to noise ratio (i.e. the proportion of fragments homologous with the probe) in the captured libraries and can decrease the numbers of homologous fragments retrieved.

**Vsearch RAD loci clustering (RAD-ref)**. High quality reads of RAD probes were clustered by similarity using <u>Vsearch</u> [32] to obtain loci for further mapping the reads from the hybridization-capture libraries. Before Vsearch run, we converted cleaned fastq files into fasta format using the <u>fasta to fastq tool</u> from the the FASTX-Toolkit package [28]. To obtain the most reliable contigs across samples, Vsearch was run in two iterations. During the first iteration, we obtained consensus clusters at the within-individual level (i.e. clustering of the raw reads for each sample independently). During the second iteration, Vsearch was run on the consensus clusters obtained from the first iteration. The second iteration allowed us to obtain consensus clusters at the among-individual level. For both iterations we ran Vsearch with various identity thresholds (0.51, 0.61, 0.71, 0.81, 0.83, 0.91, 0.93, 0.96, 0.98 for the within-individual level and 0.51, 0.61, 0.71, 0.81, 0.85, 0.88 for the among-individual level) in order to identify an optimal identity threshold for clustering, i.e. a threshold that maximizes the number of clusters with a minimal coverage of 2x and 3x, respectively. In all cases, we used the cluster_fast option for clustering. The consensus sequences of each secondary cluster were then used as locus references in subsequent alignment and SNP calling steps.

**Vsearch RAD loci clustering and extension using captured reads (RAD-ref-ext)**. To obtain RAD-ref-ext, we iteratively extended contigs of the RAD-ref using reads from the hybridization-capture library (by pooling reads contributed by all the analysed specimens) using <u>PriceTI</u> [33] with 30 cycles of extension and a minimum overlap of a sequence match to 30. The obtained references were trimmed by 60 bp at each end in order to remove sequences with putatively low-quality ends. We applied this tough threshold for RAD-ref-ext only, as probes extension can be performed on very low-coverage data, and we therefore wanted to keep the error rate (usually higher on both sequence ends) at the minimum.

**De-novo assembly from captured reads only (assembly-ref)**. Assembly was performed on the hybridization-captured reads of one good quality ethanol-preserved, sonicated, sample. Only sequences obtained from the single fresh sample were used, as stringent cleaning parameters in Trimmomatic [34], used for the reference construction, led to a large loss (up to 80%) of the reads from historical samples (and such data was therefore less optimal than that from the fresh specimen for producing a reference). Moreover, using one individual allows obtaining a more reliable reference, as any among-sample divergence can result in bubbles in the contigs and bias the assembly by oversplitting alleles. As a result we could have obtained chimeric duplicated loci presented in different contigs. We used <u>SOAP *denovo V2.04*</u> to assemble cleaned reads into contigs [35].

### Mapping and SNP calling

Reads from the hyRAD library were cleaned by Trimmomatic with milder parameters than reads used in the *de novo* assembly (keeping 60-85% of the reads in historical specimens). Read mapping was performed using <u>bowtie2-build</u> for the reference indexing and <u>bowtie2</u> for mapping [36], PCR duplicates were removed using the MarkDuplicates tool from the Picard toolkit [31] and SNPs were called with FreeBayes using the default parameters [37, 38]. To evaluate the level of DNAdamage in museum DNA samples, we used <u>mapDamage2.0</u> that rescales base quality scores of putatively post-mortem damaged bases [39] in order to minimize presence of post-mortem conversions in the resulting SNPs. Datasets for replicates were merged and analysed for SNPs with FreeBayes. Obtained VCF files were filtered by the vcffilter tool from the vcflib [40] and VCFtools [41]. In the resulting set only biallelic sites with high quality (PHRED>30), minor allele count larger than 1/6 of all, present in at least 50% of the samples and with a minimum depth of 6 were kept, indels were removed. Potential paralogs and multi-copy sites were removed based on a coverage of a standard-deviation three times higher than the mean [42]. VCF format files [43] were converted to SNP-based NEXUS files using PGDSpider converter [44] and to structure data files for every individual using the vcf-consensus tool (see also Fig 2). Updated versions of the bioinformatic pipeline can be found at https://github.com/chiasto/hyRAD.

**Fig 2.**
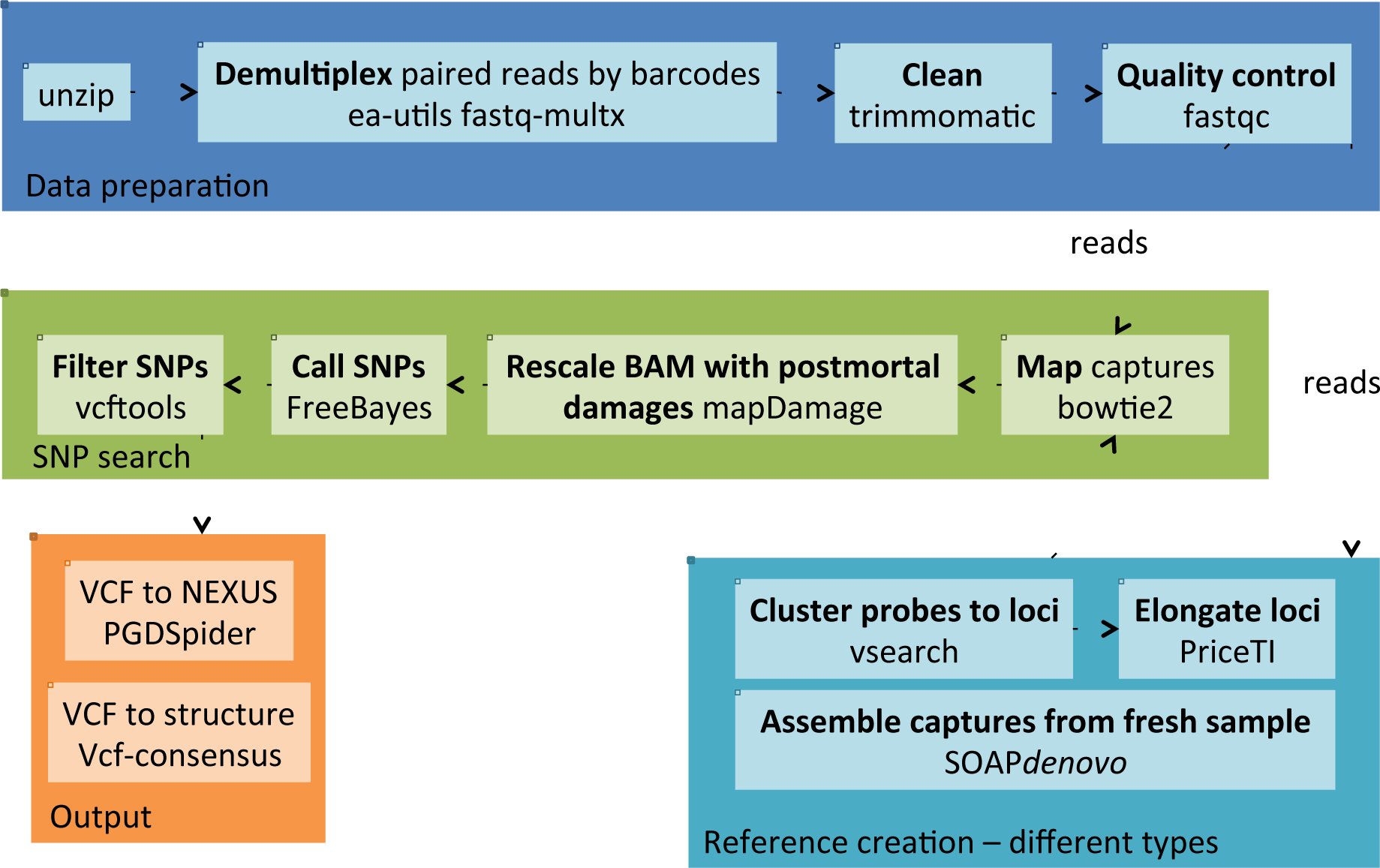
Bioinformatic pipeline used for processing hyRAD sequences. First, the reads are demultiplexed and cleaned. Different types of references were built and the captured fragments were mapped on the reference. The SNPs are then called after correcting for post-mortem DNA damages.

### Genetic structure

In order to check whether data was reflecting genetic structure, we applied fastStructure, a Structure-like algorithm adapted to large SNP genotype data [45] to the six final datasets (RAD-ref, RAD-ref-ext and assembly-ref, both for samples with and without sonication). The analyses were conducted using a *simple prior* and assuming two groups (*k*=2).

### Overlap between assembly references

To evaluate the level of overlap among the three assembly references for *L. helle* (RAD-ref, RAD-ref-ext, and assembly-ref) we used the <u>OrthoMCL</u> [46] pipeline for orthology detection. Most of the pipeline was run with the default parameters, except for Blastall and MCL clustering steps. Here, we used more stringent parameter values (e-value of 0.0001 and MCL was run with an inflation parameter of 2.0) in order to reduce chances of detecting false orthology groups. As a result, we obtained clusters of contigs being contributed by the three assembly references. We then counted how many of these clusters – presumably corresponding to homologous loci – were shared among the available reference assembly approaches. Eventually, to reveal the number of RAD loci present in the references, reads of the raw RAD library were mapped on RAD-ref and assembly-ref using bowtie2 [36] and levels of mapping were compared.

### Proof of concept: application of hyRAD to *Oedaleus decorus*

The utility of the method was further validated using 49 museum and fresh samples of a Palearctic grasshopper species for which a marked east-west spatial genetic structure was identified previously [22]. The catalog was built based on eight specimens from the captured library that showed the largest number of reads and spanned the species’ distribution area (Switzerland, Italy, Spain, Russia), using the method that yielded the highest number of contigs and produced consistent genetic structure in *L. helle* (assembly-ref, i.e., *denovo* reference built using SOAP*denovo*, see Results and Discussion section and (Table 4). The generated contigs were blasted against GenBank databases for bacterial, fungal and technical sequences with a minimum E-value threshold of 0.1. Endogeneous contigs of each samples were assembled to generate the final reference using Geneious V9.0.2 [47]. Filters were subsequently applied to keep only high quality and informative SNPs as for the *L. helle* libraries. The final VCF matrix was converted into Structure format using PGDSpider V2.0.9.0 [44] and population structure was inferred using fastStructure [45] using a *simple prior* and assuming two groups (*k*=2). Quantum GIS V2.4.0 was used to present the geographic distribution of the genetic clusters [48].

**Table 4.** Data on obtained references for *L. helle* (RAD-ref, RAD-ref-ext, assembly-ref) and *O. decorus* (assembly-ref).

| Reference | Number of contigs | Largest contig (bp) | Total length (bp) | N50 |
| --- | --- | --- | --- | --- |
| RAD-ref (*L. helle*) | 25 478 | 544 | 5 445 942 | 209 |
| RAD-ref-ext (*L. helle*) | 24 820 | 851 | 2 613 024 | 98 |
| assembly-ref (*L. helle*) | 304 161 | 2 352 | 35 579 979 | 666 |
| assembly-ref (*O. decorus*) | 408 851 | 13 103 | 119 789 911 | 321 |

## Results and Discussion

### Sequencing and data quality

***Lycaena helle* libraries**. RAD-seq libraries sequencing yielded 14,188,023 and hybridization-capture libraries 16,636,502 raw reads: 8,217,522 for sonicated and 8,418,980 for non-sonicated samples. Additional sequencing of the reference library from the ethanol-preservedspecimen (used for the RAD-ref creation) yielded 4,703,744 reads. The proportion of reads kept after quality filtering varied with sample age and preparation method. For the ethanol-preserved sample, 89.8% of reads from sonicated and 89.4% from non-sonicated sample were retained. For the 30 years old samples the mean was 74.3% and 79.6%, and for 58 years old samples 70.8% and 73.8% for sonicated and non-sonicated samples, respectively.

***Oedaleus decorus* libraries**. Hybridization-capture libraries yielded a total of 69,306,042 raw reads. After quality filtering, 80.3% of reads were kept among all the samples.

### Comparison of references obtained for *L. helle* libraries

**Proportion of single-hit alignments and SNP numbers**. Consensus clustering of the RAD-based reference within individuals produced the largest number of clusters with 2x and 3x coverage with a clustering identity threshold of 0.91, compared to other threshold values. Consensus clustering among individuals produced the best results with a clustering identity threshold of 0.71 (see S3 Fig).

The highest number, length and the total length of reference contigs were obtained using *de novo* assembly with SOAP*denovo* (assembly-ref; (Table 4). Both RAD-based assemblies produced an order of magnitude lower number of contigs. The extension performed on the obtained RAD reference followed by trimming of adapter sequences resulted in references with an average shorter length (lower N50) than the starting contigs—whereas priceTI extended a large number of probes, this did not reflect in a substantially higher average loci length because of further trimming of obtained contigs.

The highest levels of single-hit alignments for most of the samples, except the oldest ones, for both preparation methods (sonicated and not sonicated) were obtained when mapped on the sequenced RAD loci extended using PriceTI (RAD-ref-ext). This method was followed by *de novo* assembly using reads from the hybridization-capture library from a single fresh specimen (assembly-ref) and mapping on the RAD loci (RAD-ref); although the difference between the last two approaches was not large (Fig 3). Only for the oldest samples as well as in the non-sonicated fresh sample, *de novo* reference provided slightly better results.

**Fig 3.**
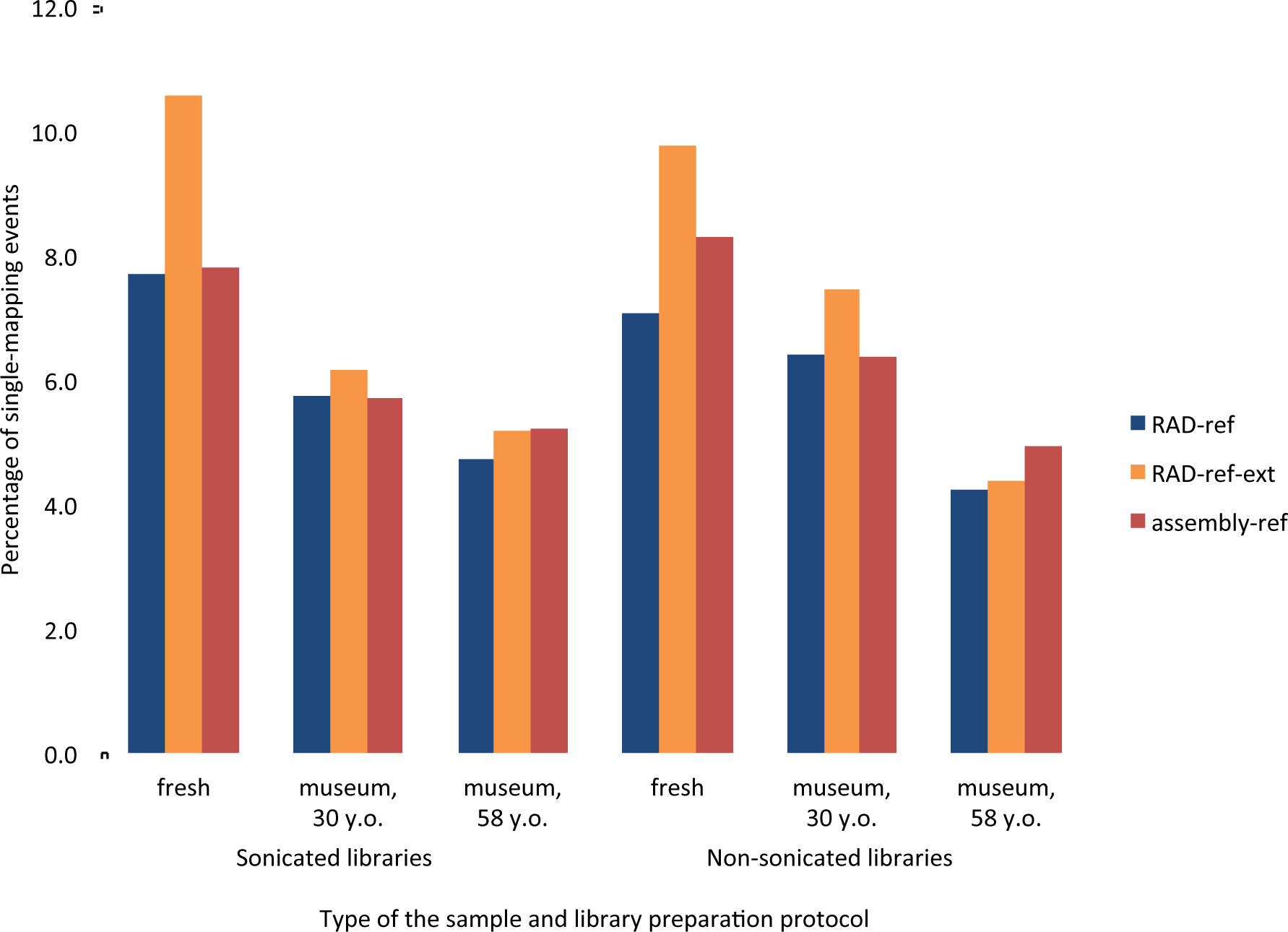
Percentage of the captured reads showing unique mapping events for different types of DNA preparations and bioinformatic pipelines.

In terms of the number of SNPs retained after coverage, paralogs and among-samples overlap filtering, the RAD-ref pipeline detected the highest numbers of loci, regardless of sample age and preparation (Fig 4). No clear correlation with sample age could be observed, although all the methods provided the lowest SNP numbers in the fresh samples – most likely an effect of small genetic distance between the reference sample and the fresh specimens (as the fresh sample was used both for the reference and the aligned sample, only heterozygote sites account for SNPs here). A higher number of SNPs was detected in the sonicated library only for the fresh specimens.

**Fig 4.**
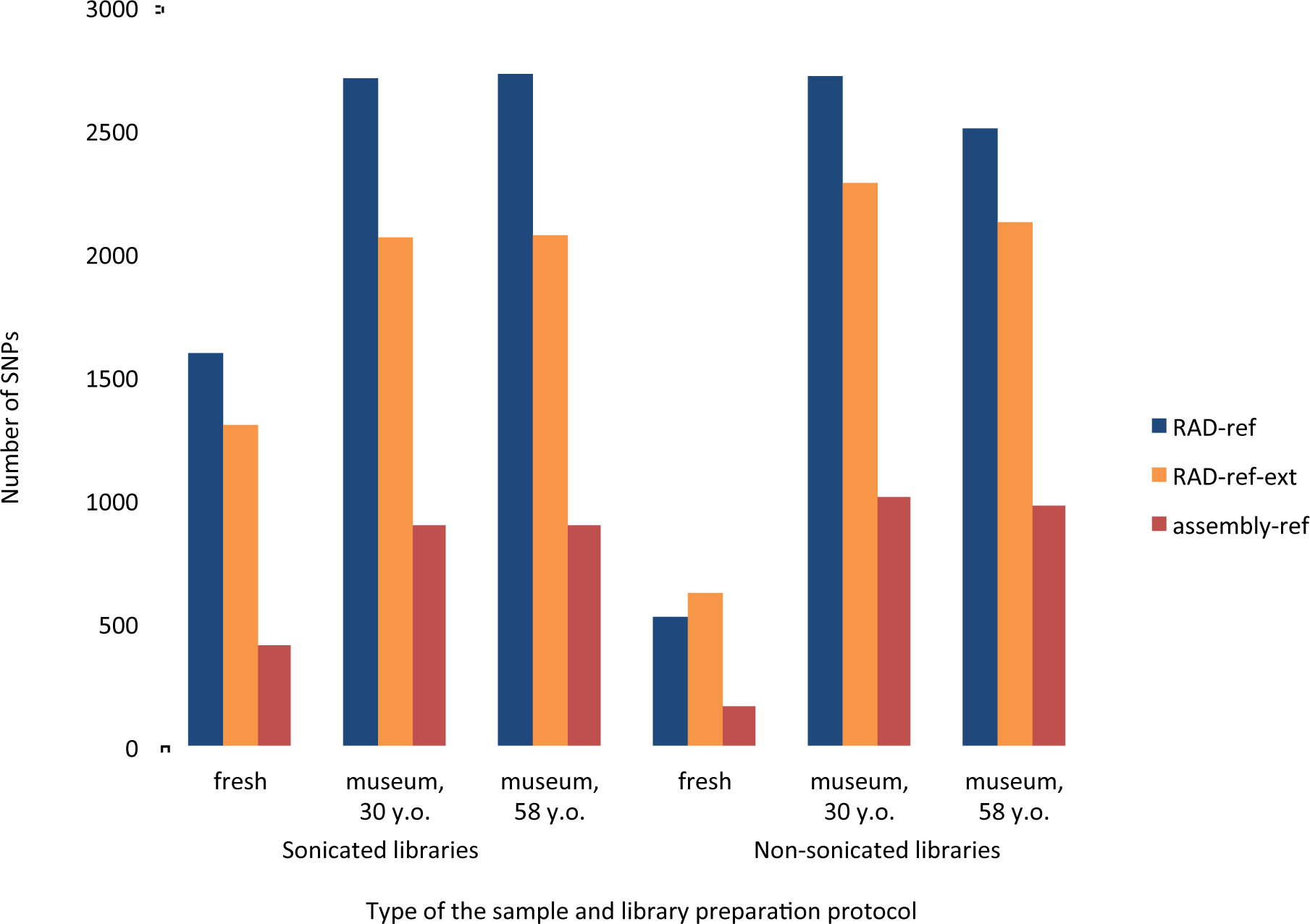
Mean number of SNPs per sample obtained for different types of DNA preparations and bioinformatic pipelines.

RAD-sequencing datasets depend on the presence of the restriction sites and therefore any polymorphism in such sites leads to either missing loci or alleles. As our method does not depend on the restriction site presence, combined with the high number of gathered SNPs, this allows obtaining largely filled data matrices. Matrix fullness was >50% in all cases:

- assembly-ref, non-sonicated: 66.1%
- assembly-ref, sonicated: 63.4%
- RAD-ref, non-sonicated: 52.0%
- RAD-ref, sonicated: 69.1%
- RAD-ref-ext, non-sonicated: 63.4%
- RAD-ref-ext, sonicated: 72.5%

Lacking an objective criterion for assessing the ‘best’ performing method for building the catalog, we tested which of the three references created the dataset producing the expected spatial division of genetic structure between Finnish and Romanian *L. helle* samples. The spatial genetic structure inferred by fastStructure revealed that the eight samples are divided into two clusters of, respectively, seven Finnish *vs.* one Romanian sample only when using the non-sonicated library mapped to the assembly-ref catalog. This result is in agreement with the hypothesis that when using sonicated DNA, we are at high risk of incorporating contaminant DNA which can blur the signal (Matthias Meyer, Max Planck Institute for Evolutionary Anthropology, Liepzig, DE; personal communication). This is an interesting result as BLAST analyses on the six catalogs did not retrieve differences in the level of known contaminants (see below).

**Loci overlap among the assembly methods**. About 15.2% of the reference loci obtained via *de novo* assembly were shared with those based on the clustering of RAD-seq reads (either RAD-ref or RAD-ref-ext; (Fig 5). In contrast, an appreciable fraction of the obtained reference loci (59.7%) were unique to the assembly-ref approach. When mapping raw RAD reads on RAD-ref, 26.3% did not map, 42.4% mapped once and 31.3% mapped more than once. This shows that RAD-ref summarizes ca.% ¾ of the reads from the RAD dataset. In contrast, when mapping raw RAD reads on assembly-ref, 78,3% of reads did not map, 21,6% mapped once and 0,1% mapped more than once. This result shows that assembly-ref possibly contains three quarters of all loci that are not homologous to the RAD probes. Such low signal to noise ratio (targeted reads to the number of total reads) is most likely a result of background carryover in the hybridization capture step, a phenomenon which can have many sources., It could result from ‘daisy-chaining’ of the captured fragments [49, 50], where partially complementary DNA molecules hybridize with the other fragments that are already hybridized to the probes. We can however discard this explanation as a primary reason for the background carryover as extending of RAD probes did not produce longer contigs (in RAD-ref-ext assemblies). Another likely reason could be carryover of random DNA fragments with repetitive sequences. The extent of such process can be significantly reduced by adding blocking agents to the hybridization mix (typically Cot-1 as was performed for the *O. decorus* dataset [51] or salmon sperm DNA), as well as optimizing hybridization temperature and wash stringency to increase capture efficiency. Such developments are desirable because they increase the percentage of reads matching the loci of interest and eventually improve the overall sequencing coverage.

**Fig 5.**
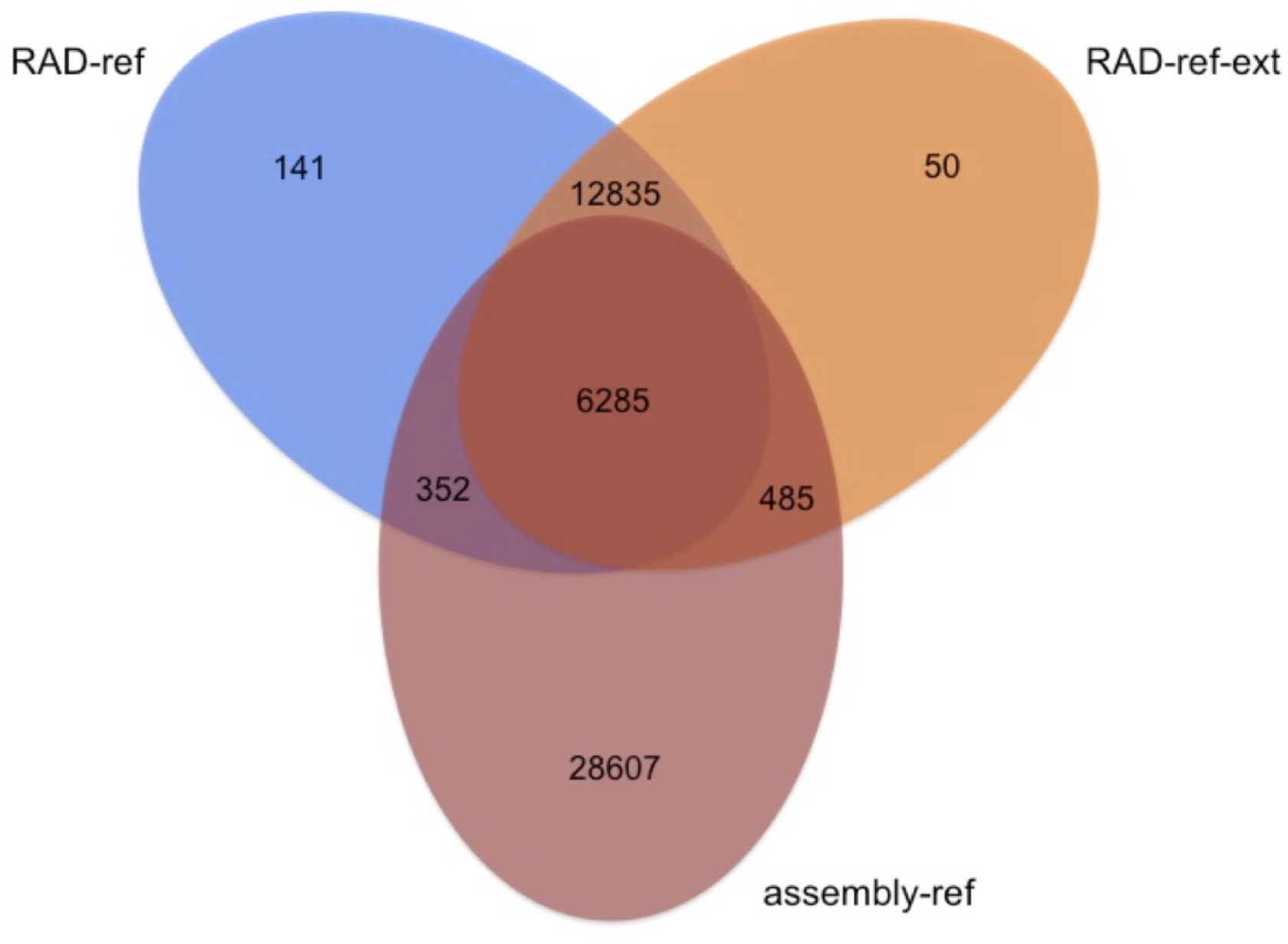
Number of loci obtained using different bioinformatic approaches, identified using the OrthoMCL [42] pipeline for orthology detection.

**Effects of sample preparation and age on the numbers of SNPs obtained and the exogeneous DNA content**

Reads can be mapped on a reference either once with a highest score (i.e., single mapping) or on more than one region of reference with close scores (i.e., multi mapping). The reasons for multi-mapping events can be biological (e.g., paralog sequences) or technical (splitting single loci into more reference loci), nevertheless these mapping events cannot be used for SNP calling and offer another benchmark for the assembly methods used. Differences in the number of single mapping events and in the numbers of SNPs obtained were not substantial between sonicated and non-sonicated samples, and depended on the sample age and the bioinformatic pipeline used (Figs 3 and 4). We expected that museum specimens should perform better without sonication, as the DNA was already visibly fragmented, and sonication of museum specimens may increase the levels of exogenous DNA contamination (by fragmenting intact fungal or bacterial DNA contaminating museum samples; M. Meyer, pers. comm.). In terms of mapping events, whereas the fresh sample usually showed a better ratio of single-to multi-mapping events when sonicated, there was a trend towards a higher percentage of unique mapping events for non-sonicated 30-years old museum specimens, on average 1.3 times higher, irrespective of the reference used (the difference was less clear for 58-years old museum specimens, and depended on the reference used). We would therefore advise not to sonicate DNA obtained from the museum specimens, which significantly cuts down the price and time required for library preparation, except in cases when no signs of degradation are observable on the DNA profile. As levels of DNA degradation of contemporary samples may vary, one may consider that the sonication step should be advisable when working with well-preserved DNA. However, this is still an open question, as whereas BLAST searches did not retrieve higher fractions of contaminants in sonicated *vs.* non-sonicated libraries, the expected population structure was retrieved was the non-sonicated one mapped on assembly-ref.

As one of the main types of post-mortem DNA degradation is deamination of cytosines, highly damaged ancient or museum DNA samples are usually characterized by higher uracil content [52-54]. In classical library preparation protocols, the usage of a proofreading polymerases should stall the chain elongation in the presence of uracil and thus reduce the misincorporation errors in the final dataset. On the other hand this approach might not be optimal for highly degraded ancient DNA samples, where a large fraction of DNA fragments may carry cytosine to uracil misincorporations [53]. Moreover the usage of a proofreading polymerase does not prevent the misincorporations caused by direct deamination of methylated cytosine to thymine, or less common deamination of guanidine to adenine [52, 54]. In the protocol used above [15], the second strand synthesis was performed using Klenow Fragment (3’ → 5’ exo-), lacking proofreading ability, and thus approximately half of the resulting DNA fragments should have cytosine to uracil misincorporation substituted for thymine, amplifiable by proofreading DNA polymerase. We thus opted for a bioinformatic post-processing way of filtering-out such bases. Post-mortem damage in the sequenced samples was assessed by mapDamage2.0, which rescales sequence files by downscaling quality scores of likely post-mortem damaged bases. As some SNPs became filtered by lower quality scores after the rescaling, the number of SNPs is decreased after mapDamage2.0. We expected higher number of discarded SNPs in the oldest samples, because of a higher proportion of DNA damage occurring with time. The proportion of SNPs discarded after applying mapDamage2.0 was the highest among the 58 years old samples (1.46% for RAD-ref; 1.89% for RAD-ref-ext; 2.36% for assembly-ref), although relatively low, given the samples’ age and preservation type (see S1 Table).

It is worth mentioning, that the higher number of SNPs detected in libraries from museum specimens, comparing to the fresh samples, is not an effect of post-mortem damage (an opposite trend was detected with the higest proportion of type II transitions to transitions [55] and transversions present in the fresh specimens; S4 Fig)

### Application: spatial genetic structure of *Oedaleus decorus*

The *de novo* reference catalog was composed of 408,851 reference contigs. The N50 length was 321 bp and the total length was 119,789,911 bp. Among the total number of contigs, 9% were shown to be of exogeneous origin by the BLAST search, either against fungi and bacteria GenBank databases or against technical sequences. Such a level of contaminants is expected here, as in contrast to the *L. helle* references, which were built from fresh samples, the *O. decorus* assembly-ref was based on eight specimens from the captured library—either fresh or pin—mounted—that showed the largest number of reads. A total of 4,783,774 informative sites were retrieved after SNP calling. After removing indels and low quality sites, 125,890 sites were conserved. Keeping only biallelic loci with a minor allele count of at least 6, with data fullness higher or equal to 50% of the samples, we obtained 6,046 loci. Finally, we conserved 2,979 SNPs after the removal of potential paralogous sites. The median depth for each SNP was 10. On average, each of the 49 samples were characterized by 1864 SNPs and each SNP was found in 32 individuals (62.7% of matrix fullness).

The spatial genetic structure inferred by fastStructure revealed two geographically distinct clades in the west and the east of the Palearctic (Fig 6). This result supports the eastern-western split previously highlighted in *Oedaleus decorus* based on mtDNA amplicons [22]. This demonstrates that hyRAD is a reliable technique to infer spatial genetic structure from both fresh samples and museum samples collected at various time points in the past.

**Fig 6.**
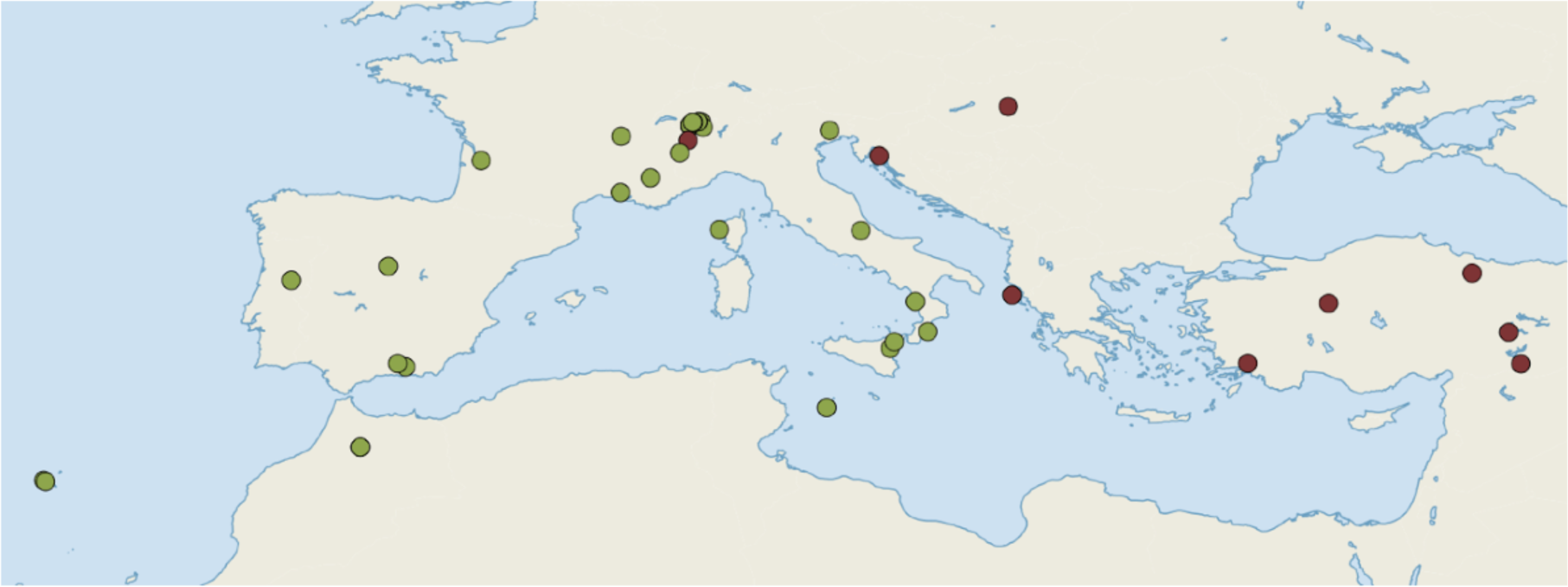
Spatial genetic structure of *O. decorus* inferred using fastStructure with *k*=2 and simple priors. Colours denote the two different genetic groups supported by a previous study relying on mtDNA markers [22].

## Conclusions

Here, we present a method for obtaining large sets of homologous loci from museum specimens, without any a priori genome information. Despite the differences in single-mapping events among samples of different ages were retrieved, the obtained numbers of SNPs were not significantly different for the two age classes of the *L. helle* museum specimens. We obtained around a thousand of SNPs from *L. helle* samples up to 58 years old, confirming that it can be successfully applied in the field of museum genomics. The application of the catalog-building method that was the most promising for resolving population genetic structure (i.e., non-sonicated library mapped to assembly-ref) to the grasshopper *O. decorus*, confirmed the usefulness of hyRAD to retrieve phylogeographic data using museum samples up to one hundred years old. Our method does not require time-consuming and costly probes design and synthesis, nor access to fresh samples for RNA extraction, making it one of the simplest and most straightforward technique for obtaining orthologous loci from degraded museum samples.

In the protocol, we applied a modified shotgun library preparation method, optimized for degraded DNA from museum specimens [15]. However, the capture protocol presented here can be applied to any type of library preparation, including commercial ones, simplifying the workflow and cutting down the preparation time.

We also explored several bioinformatic approaches for loci assembly from the captured libraries, a crucial step when working on organisms without a reference genome. Identifying the most appropriate catalog-building method may depend on the goals of each study. In our case, the pipeline that was the best at identifying population structure in the butterfly was relying on a non-sonicated library mapped to the *de novo* reference assembly from captured reads from a single ethanol-preserved specimen, using SOAP*denovo* assembler (assembly-ref). Despite the fact that a maximum of 26% of the obtained sequences mapped to the references and the proportion of single mapping events were not higher than 10% on average (Fig 3), we could successfully call around a thousand of loci in each case (Fig 4), with high coverage across the samples.

Importantly from the wetlab protocol perspective, in the hybridization step, we have used blocking oligonucleotides to prevent ‘daisy-chaining’ of captured sequences by adapter sequences’ homology. Using a blocking agent preventing similar chaining caused by repetitive sequences (Cot-1 DNA, applied to the grasshopper libraries, see below [51]) as well as optimizing the conditions of hybridization and capture reactions for increased stringency (e.g., by decreasing hybridization temperature and the stringency of the washes using higher concentration of SDS and/or lower concentration of SSC) may further increase the hybridization efficiency and thus the numbers of reads mapping on the reference and reduce the number of low-coverage loci.

The *de novo* assembly building pipeline produced the largest contig of 2,352 bp for the butterfly and 13,103 bp for the grasshopper dataset. Although the mean length of the assembled contigs was much smaller, our method also allows retrieving longer sequences than the length of the probes used. The reason for this is that captured sequences hybridize with other DNA fragments with homologous sequences, flanking the probe sequence (i.e., ‘daisy-chaining’ [49, 50]). This may lead to enrichment across larger fractions of genome, a side effect of our method, that can be utilized for assembly of larger contigs by using longer probes and capturing longer targets.

The method presented here, although based on the restriction enzyme digestion of DNA to create the random genomic probes, does not depend on the restriction site presence in the captured library. This represents a significant improvement over classical RAD-sequencing datasets, in which increase in the phylogenetic distance among samples is correlated with an increase in the number of missing sites [56-61], sometimes leading to conflicting signals between RAD-and capture-based datasets [62], or are characterized by the presence of null alleles that lead to heterozygosity or *F*_ST_ underestimation [9, 10]. In this aspect, our approach is similar to other capture-enrichment protocols, such as UltraConserved Elements [5] or exome-capture [4], with the benefit of much simpler and less expensive probe generation, without access to genome information or fresh specimens for RNA isolation. Not relying on the presence of restriction site, the method presented here should be also useful for broader phylogenetic scales, allowing sequencing homologous loci from more divergent taxa, which would not be possible to retrieve using classical RAD-seq approaches.

## Acknowledgments

We thank Alan Brelsford, Alicia Mastretta-Yanes and Pawel Rosikiewicz for their help with developing RAD-sequencing protocols. Jairo Patiño tested early versions of the protocol and provided valuable feedback. Roger Vila and Gerald Heckel kindly provided fresh samples for the study. We thank the following museum curators for providing collection samples: Hannes Baur (Natural History Museum, Bern, Switzerland), Anne Freitag (Zoological Museum, Lausanne, Switzerland), Rod Eastwood (ETH Entomological Collection, Zurich, Switzerland), Daniel Burckhardt (Natural History Museum, Basel, Switzerland), Peter Schwendinger (Natural History Museum, Geneva, Switzerland), Barabara Oberholzer (Natural History Museum, Zurich, Switzerland), George Beccaloni (Natural History Museum, London, UK), Lauri Kaila (Finnish Museum of Natural History, Helsinki, Finland). We also thank Brent Emerson for his constant support during the development of this method.

## Supporting Information

S1 Fig. Profile of the RAD-probes precursor, the RAD-seq library. Left panel, *X*-axis: fragment size (semi-*log* scale); *Y*-axis: fragment density (Relative Fluorescent Units). Right panel, gel-like representation of the left panel.

S2 Fig. Profile of the re-amplified capture library after AMPure purification. Left panel, *X*-axis: fragment size (semi-*log* scale); *Y*-axis: fragment density (Relative Fluorescent Units). Right panel, gel-like representation of the left panel.

S3 Fig. Illustration of the clustering optimization of RAD-ref assembly clustering thresholds using Vsearch. *X*-axis: clustering threshold; *Y*-axis: number of clusters with 2x (red) or 3x (green line) coverage. The top panel shows within-sample, whereas the bottom panel shows among-sample clustering results. The optimal threshold optimizes the number of the clusters with 2x and 3x coverage.

S4 Fig. Proportion of type II transitions to all the transitions and transversions for each of the reference catalog, with and without post-mortem bias correction. The data shown are for the fresh sample with DNA sonication and the museum samples without sonication. The left plot shows values without and the right plot with mapDamage2.0 correction.

S1 Table. mapDamage2.0 results based on each of the three reference catalogs for *L. helle* analyses, with the number of obtained SNPs (with and without application of mapDamage2.0).

